# Balancing the influenza neuraminidase and hemagglutinin responses by exchanging the vaccine virus backbone

**DOI:** 10.1101/2020.12.08.416099

**Authors:** Jin Gao, Hongquan Wan, Xing Li, Mira Rakic Martinez, Yamei Gao, Zhiping Ye, Robert Daniels

## Abstract

Virions are a common antigen source for many viral vaccines. One limitation to using virions is that the antigen abundance is determined by the content of each protein in the virus. This caveat especially applies to viral-based influenza vaccines where the low abundance of the neuraminidase (NA) surface antigen remains a bottleneck for improving the NA antibody response. Here, we used the WHO recommended H1N1 antigens (2019-2020 season) in a systematic analysis to demonstrate that the NA to hemagglutinin (HA) ratio in virions can be improved by exchanging the viral backbone internal genes, thereby preserving the antigens. The purified inactivated virions with higher NA content show a more spherical morphology, a different balance between the HA receptor binding and NA receptor release functions, and induce a better NA inhibitory antibody response in mice. These results indicate that influenza viruses support a range of ratios for a given NA and HA pair that can be utilized to produce viral-based influenza vaccines with increased NA content which can elicit more balanced inhibitory antibody responses to NA and HA.

## Introduction

Virions are the extracellular infectious form of a virus. In general, they consist of the viral genome surrounded by an outer protein shell, or a viral protein-containing envelope (membrane). The main function of the virion surface proteins is to initiate the infection process by mediating specific interactions with the host cell, which is crucial for viral genome delivery. These essential functions and the accessibility of the viral surface proteins are one of the primary reasons why many of them are also effective vaccine antigens. Accordingly, virions, which possess high surface protein content and are relatively easy to isolate, are commonly used as an antigen source for many vaccines including those against polio, hepatitis A and influenza viruses [1]. While this approach has been effective, it can also overlook protective surface antigens that are not abundant in virions.

Influenza virions possess two major surface antigens, hemagglutinin (HA or H) and neuraminidase (NA or N), that are both capable of eliciting a protective antibody response against influenza virus infections [2–5]. However, the HA amounts in the viral envelope are generally much higher than the NA amounts and can differ between influenza virus strains [6–8]. One contributing factor to the HA to NA ratio in a virion is the requirement to balance the low affinity receptor binding function of HA with the receptor removal function of NA [9–12]. This balance requirement continues to complicate the inclusion of NA in viral-based vaccines, as the ratio in virions is likely a unique trait for each HA and NA pair and current influenza vaccines are recommended to contain four different pairs of HA and NA, two from influenza A viruses (IAVs) and two from influenza B viruses [13].

The two IAV antigen pairs are from the H1N1 and H3N2 subtypes and the representative field strains are chosen yearly by the WHO based on surveillance data combined with serological and sequence analysis of the HA antigen. The most common influenza vaccines are derived from inactivated virions that are produced in eggs, or more recently in cells [13]. To optimize the vaccine yields, manufacturers generally use influenza candidate vaccine viruses (CVVs), which are generated by reassortment of the representative HA and NA antigens with a non-pathogenic, high-growth influenza genome backbone such as the one from the A/PR8/34 (PR8) strain [14]. Despite containing NA, CVVs are benchmarked against the wild-type strain based on HA content, potentially resulting in the selection of strains that possess low NA amounts. While the underlying cause remains unclear, numerous studies have shown that influenza vaccines elicit a suboptimal NA antibody response [15–17], leading to the speculation that HA is immunodominant, or that NA amounts in the vaccine are insufficient or of poor quality.

The IAV genome is comprised of eight single-stranded influenza RNA gene segments, which encode for one or more viral proteins [18]. Several studies have used reverse genetics to change the virion incorporation of HA or NA by altering the promoter region, codons, or N-linked glycosylation sites in their respective gene segment [7, 19–21]. Similar investigations have found that exchanging the internal gene segments can also affect the NA and HA amounts in virions [22–25], a trait supported by more mechanistic analysis of NA and HA expression in cells [26, 27]. Here, we investigated if the PR8 influenza vaccine virus backbone can be modified to increase the NA virion content and balance the HA and NA antibody responses without changing either antigen. In addition to the HA gene segment, our results demonstrate that the NA virion content is also dependent on the six internal influenza gene segments or the virus backbone. The properties corresponding with the increased NA virion content are presented along with data showing that the inactivated virions elicit a stronger NA antibody response in mice without an associated loss in the HA antibody response. The mechanistic and translational implications of these results for producing viral-based vaccines that elicit more balanced NA and HA antibody responses are discussed.

## Results

### NA production by H1N1 influenza viruses in MDCK cells and eggs is strain dependent

During previous studies, we observed that single-gene reassortant IAVs using NAs from subtype 1 (N1) are more easily rescued with a backbone from the A/WSN/33 (WSN) strain compared to others and produce high NA levels [19, 28, 29]. To investigate this observation in a more controlled manner, single-gene reassortant IAVs carrying N1 antigens form the WHO recommended CVVs A/California/07/2009 (N1-CA09) or A/Brisbane/02/2018 (N1-BR18) were rescued using the seven gene segment backbone from WSN and PR8 (Fig. 1A). PR8 was chosen for comparison because it is highly egg-adapted, like WSN, and often used to generate CVVs with high HA yields [14, 30]. Initially, MDCK cells were infected with the four viruses at equal MOIs and the NA amount produced by each virus was analysed when the infection reached completion (CPE was ~100%). For each NA, the viruses with the WSN backbone displayed higher NA sialidase activity in the virus-containing cell medium and in the infected cell remains, indicating higher NA amounts were produced (Fig. 1B, top panel). In contrast, both viruses with the PR8 backbone generated higher hemagglutination unit (HAU) titres in the cell medium (Fig. 1B, bottom panel). Similar results were obtained using embryonated eggs, as the virus with the WSN^N1-BR18^ virus produced higher NA levels in the allantoic fluid and lower HAU titres than PR8^N1-BR18^ virus (Fig. 1C), indicating the NA differences are likely due to strain specific viral gene products.

**Figure 1.**
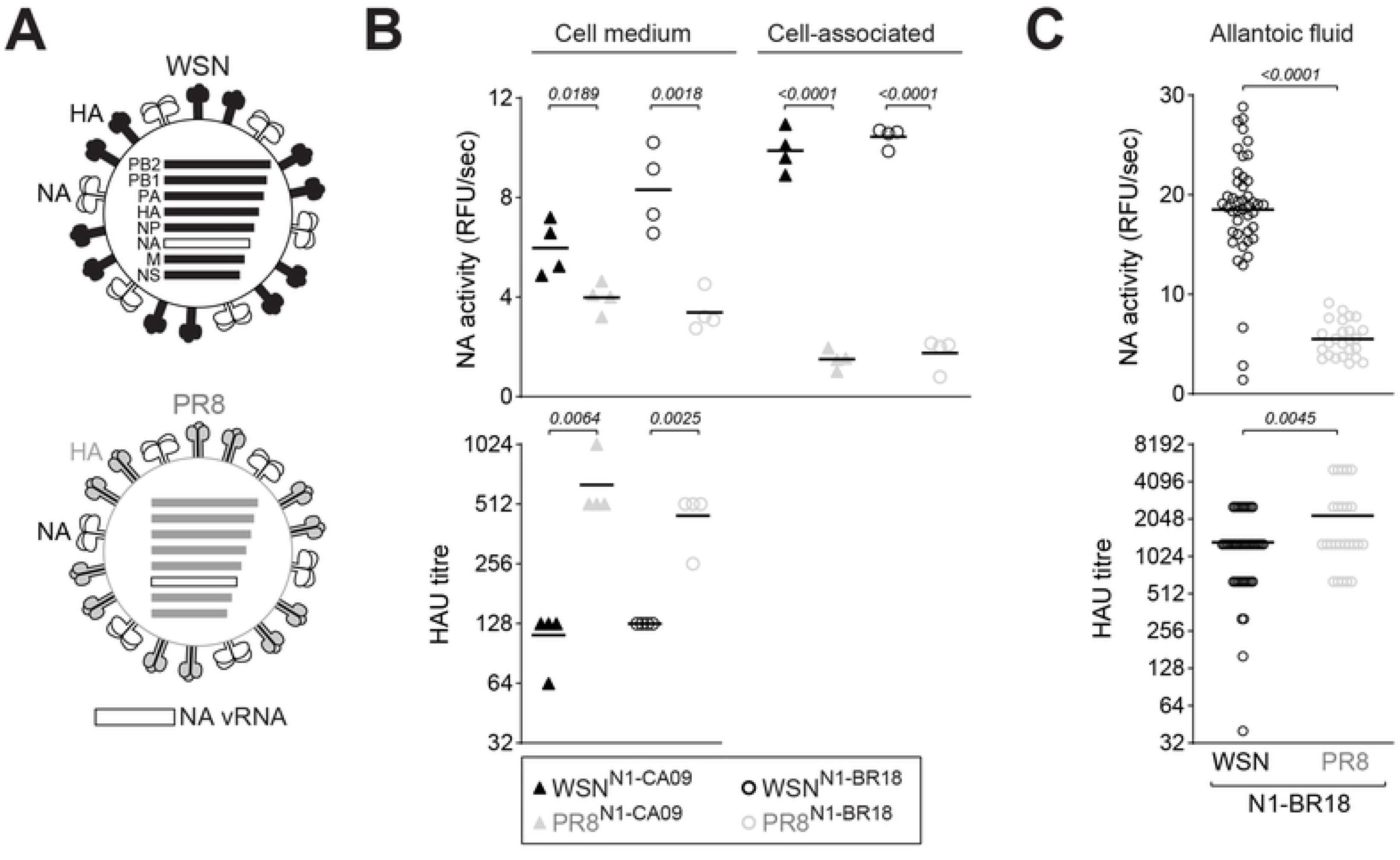
NA production by single-gene IAV reassortants is backbone-dependent. **A.** Diagram of the NA single-gene reassortant IAVs with a WSN and PR8 backbone. Viral RNA (vRNA) gene segments unique to each virus are color coded, including the HA surface antigen. **B.** The NA activity and HAU titres were measured from MDCK cells infected with the indicated viruses at a MOI of ~0.001. Upon completion of the infection, the medium and cell remains were collected, separated by centrifugation, and analyzed using equal sample amounts. The data from 4 independent biological replicates is displayed with the mean (bar) and the *P* values calculated from a two-tailed unpaired t-test, 95% CI. **C.** Embryonated eggs were infected with the indicated viruses for three days prior to harvesting and measuring the HAU titers and NA activity in equal volumes of allantoic fluid. The individual egg data from three independent experiments with different batches of eggs is shown with the mean (bar). The *P* values were calculated from a two-tailed unpaired t-test, 95% CI.

### Higher NA activity levels correlate with increased virion incorporation of NA

To investigate if the increased NA levels reflect higher virion incorporation, we set out to purify inactivated WSN^N1-BR18^ and PR8^N1-BR18^ viruses from eggs. CVVs for egg-based vaccines are commonly inactivated with either beta-propiolactone (BPL) or formaldehyde (FA) [1, 31], which can modify viral proteins in addition to ribonucleic acids [32–35]. Therefore, we first examined if BPL and FA affect the viral NA activity or HAU titre. For both viruses, BPL and FA caused similar concentration-dependent decreases in allantoic fluid NA activity with FA resulting in slightly more variability (Fig. 2A). In contrast, HAU titres significantly decreased with high concentrations of BPL (Fig. 2B), especially WSN^N1-BR18^, suggesting this virus is more susceptible to BPL modification or contains less HA. Following confirmation of viral inactivation by three passages in eggs, we proceeded with 0.05% BPL because it showed little impact on NA activity and HAU titers, has a short half-life, and reacts with water to produce a non-toxic compound [32].

**Figure 2.**
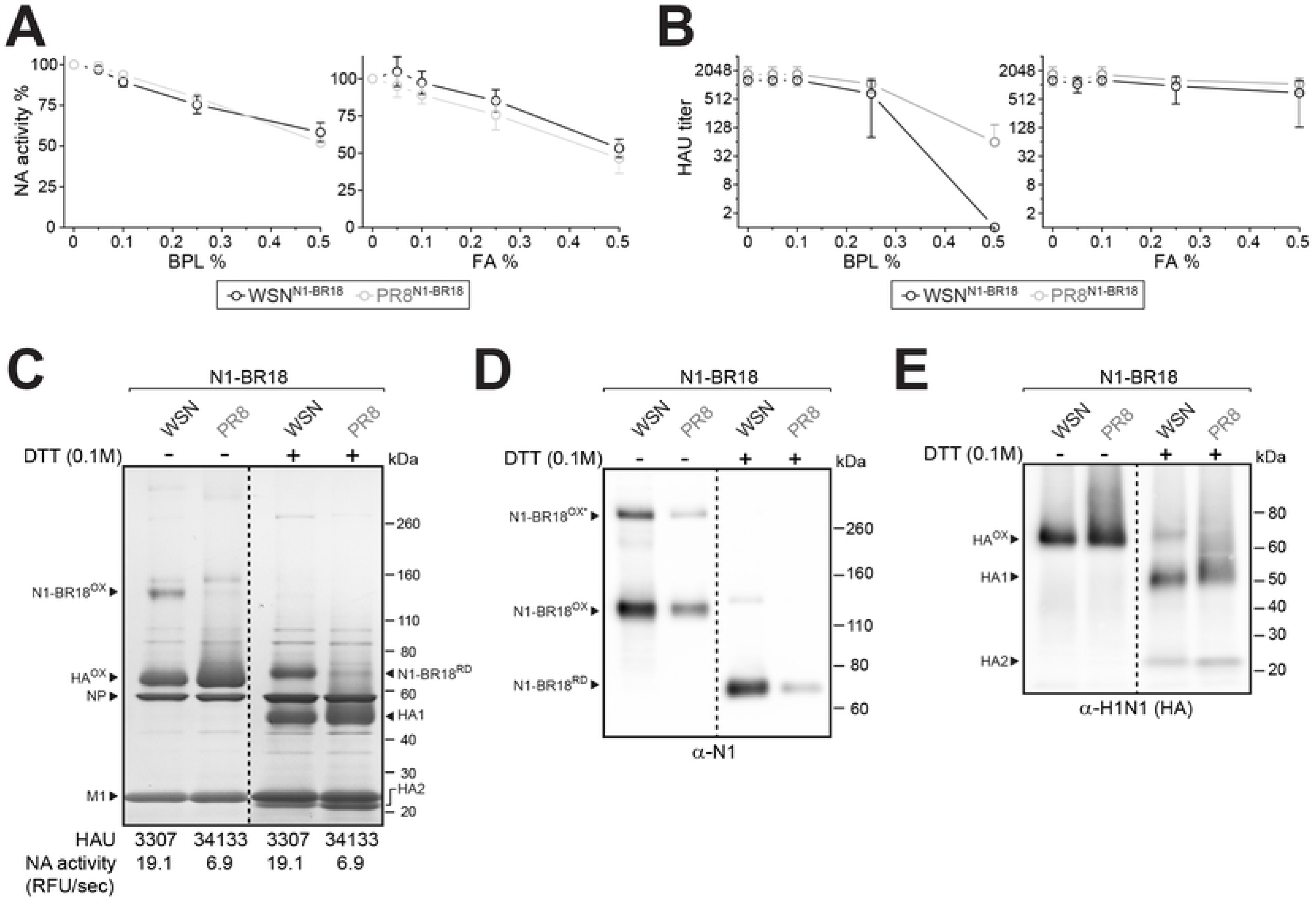
Egg produced WSNN1-BR18 virions incorporate more NA than PR8N1-BR18 virions. **A** and **B.** Graphs showing viral NA activity (**A**) and HAU titers (**B**) after treatment with increasing concentrations of β-propiolactone (BPL) and formaldehyde (FA). The indicated viruses in allantoic fluid were treated overnight at 4°C. NA activity of the untreated sample was set to 100%. The data from 3 independent biological replicates are displayed as the mean (circle) with the SD (error bars). **C.** Representative Coomassie stained gel of the purified BPL inactivated virions (5 μg per lane), untreated or reduced with DTT prior to resolution on a 4-12% SDS-PAGE gel. Oxidized (OX) and reduced (RD) forms of NA (N1-BR18) and HA are indicated with the viral proteins NP and M1. Note HA resolves as 2 polypeptides (HA1 and HA2) after reduction (+DTT). HAU titres and NA activity values are means from 3 independently purified batches with equal total protein concentrations (1 mg/ml). **D** and **E.** Representative NA (**D**) and HA (**E**) immunoblots of the purified virions (0.5 μg per lane), untreated or reduced with DTT prior to resolution by SDS-PAGE. Bands corresponding to oxidized and reduced NA and HA are indicated. (**D**) Asterisk indicates SDS resistant NA homo-tetramers that resolve under non-reducing conditions. (**E**) Note the H1N1 antisera has some reactivity towards NA.

After BPL inactivation and purification, the viruses were adjusted to equal total protein concentrations and analysed. On both Coomassie stained SDS-PAGE gels (Fig. 2C) and immunoblots (Fig. 2D), bands at the expected molecular weight of oxidized NA (dimers) and reduced NA (monomers) [19, 36] were more apparent for WSN^N1-BR18^ than PR8^N1-BR18^, indicating WSN^N1-BR18^ virions incorporate more NA. The increase in NA levels correlated with a decrease in HA content but did not affect the efficiency of HA processing to HA1 and HA2 or the content of the viral nucleoprotein (NP) or matrix (M1) protein (Figs. 2C and 2E). Supporting these observations, WSN^N1-BR18^ virions possessed ~3 times more NA activity, but ~10 times lower HAU titres than the PR8^N1-BR18^ virions (Fig. 2C). These data indicate that one or more WSN backbone gene product supports higher viral incorporation of recent N1s at the expense of reduced HA levels.

### HA and the backbone can influence NA virion incorporation

The two single-gene reassortant viruses possess backbones (WSN and PR8) that encode for distinct HAs raising the question of whether the observed increase in the virion NA content is dependent on HA due to potential changes in the functional balance. Therefore, we generated two double-gene reassortant viruses where the HA genes were swapped while the NA (N1-BR18) remained identical (Fig. 3A). In eggs, the virus with the WSN backbone and the HA from PR8 (WSN^H1-PR8/N1-BR18^) produced higher NA levels and slightly better HAU titres than the virus with the HA from WSN and the PR8 backbone (Fig. 3B). Following purification, bands corresponding to oxidized and reduced NA were apparent for both viruses on Coomassie stained gels (Fig. 3C) and immunoblots (Fig. 3D), which confirmed the NA levels were slightly higher in the WSN^H1-PR8/N1-BR18^ virions. The virion HA content was rather consistent, but somewhat lower M1 levels were observed in the WSN^H1-PR8/N1-BR18^ virions. In line with the Coomassie gel, WSN ^H1-PR8/N1-BR18^ virions possessed ~15% more NA activity than PR8^H1-WSN/N1-BR18^ virions, whereas the HAU titers were more than double (Fig. 3C). These results implied that HA and one or more WSN backbone gene products support increased viral incorporation of recent N1s and that the HA from PR8 generates higher HAU titres with turkey red blood cells than the HA from WSN.

**Figure 3.**
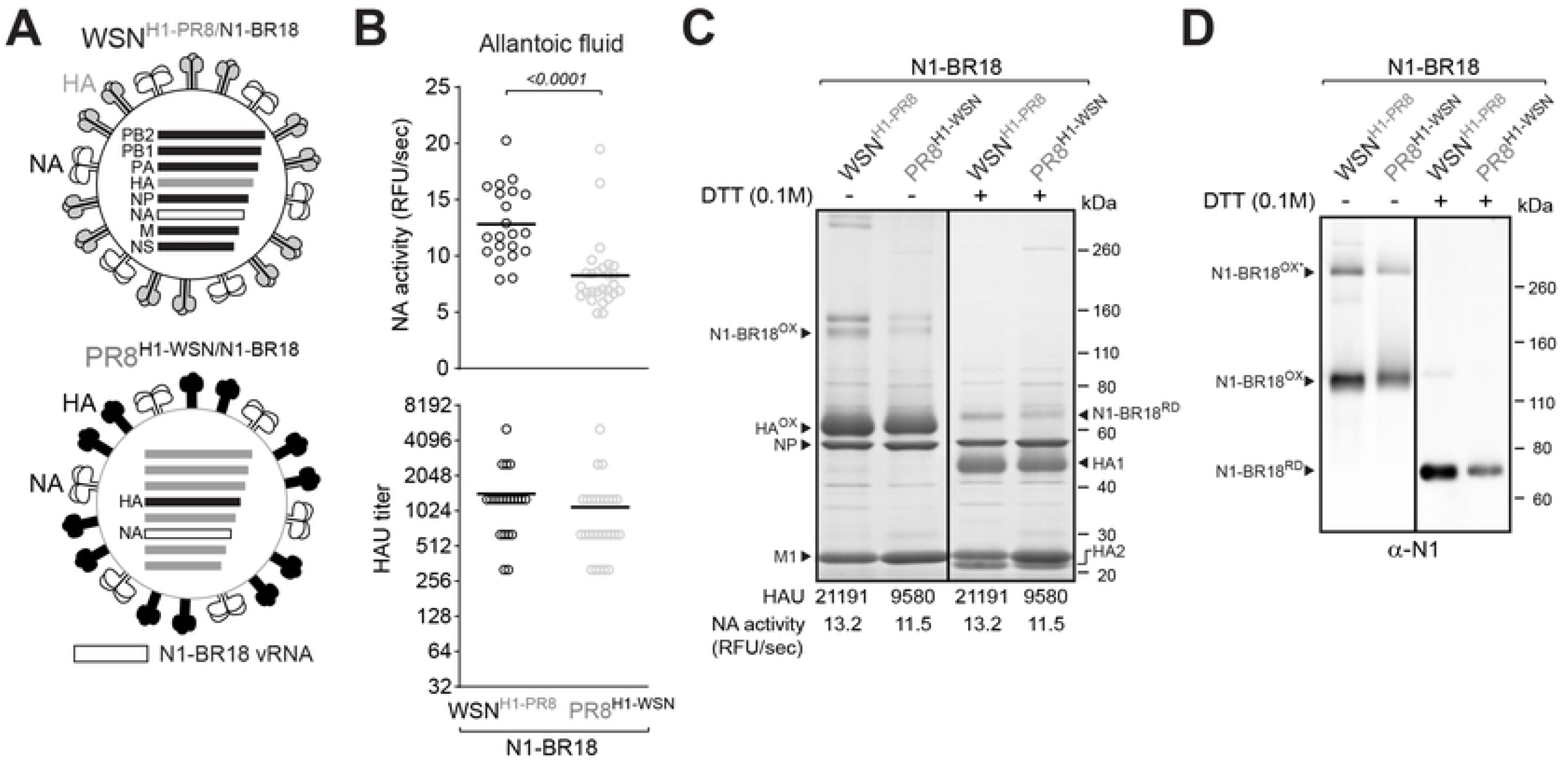
HA and the viral backbone can influence the NA virion incorporation. **A.** Diagram of the two HA and NA double-gene reassortant IAVs. The NA gene segment (N1-BR18) is the same in both viruses, but the HA gene segments from WSN and PR8 were swapped. **B.** Eggs infected with the indicated viruses was harvested after 3 days and the HAU titers and NA activity was measured using equal volume of allantoic fluid from each egg. The individual egg data from three independent experiments with different batches of eggs is shown with the mean (bar). Only significant *P* values calculated from a two-tailed unpaired t-test (95% CI) are displayed. **C.** Coomassie stained gel showing 5 μg of purified BPL inactivated WSN^H1-PR8/N1-BR18^ and PR8^H1-WSN/N1-BR18^ virions that were untreated or reduced with DTT prior to resolution on a 4-12% SDS-PAGE gel. Oxidized (OX) and reduced (RD) forms of N1-BR18 and HA are indicated along with the viral proteins NP and M1. The HAU titres and NA activity values are means from the 3 independently purified virus batches with equal total protein concentrations (1 mg/ml). **D.** NA immunoblots of the BPL inactivated purified virions (0.5 μg per lane) that were treated with DTT as indicated prior to SDS-PAGE. Bands corresponding to oxidized and reduced N1-BR18 are indicated. The asterisk denotes SDS resistant N1-BR18 homo-tetramers.

### The six internal WSN backbone gene segments support high viral incorporation of recent N1s

To mitigate the HA influence on the higher NA content in WSN virions, two double gene-reassortant viruses were generated using the WHO recommended H1 and N1 antigens for the 2019-2020 influenza season (Fig. 4A). Again, the virus with the WSN backbone (WSN^H1N1-BR18^) produced higher NA levels in egg allantoic fluid than the PR8 backbone virus (PR8^H1N1-BR18^) and rather similar HAU titres (Fig. 4B). Coomassie stained SDS-PAGE gels and activity measurements of the purified virions both confirmed that the NA levels were significantly higher in WSN^H1N1-BR18^ (Fig. 4C). Although the HAU measurements were identical, the NA increase coincided with a decrease in HA that was observed both on Coomassie stained gels and immunoblots (Fig. 4C and 4D). Using an unrelated HA subtype, H6 from the strain A/turkey/Mass/3740/1965, the virus with the WSN backbone was also observed to produce higher NA activity in the allantoic fluid (Fig. 4E) and had higher NA content (Fig. 4F), demonstrating that the WSN backbone supports high N1 virion incorporation.

**Figure 4.**
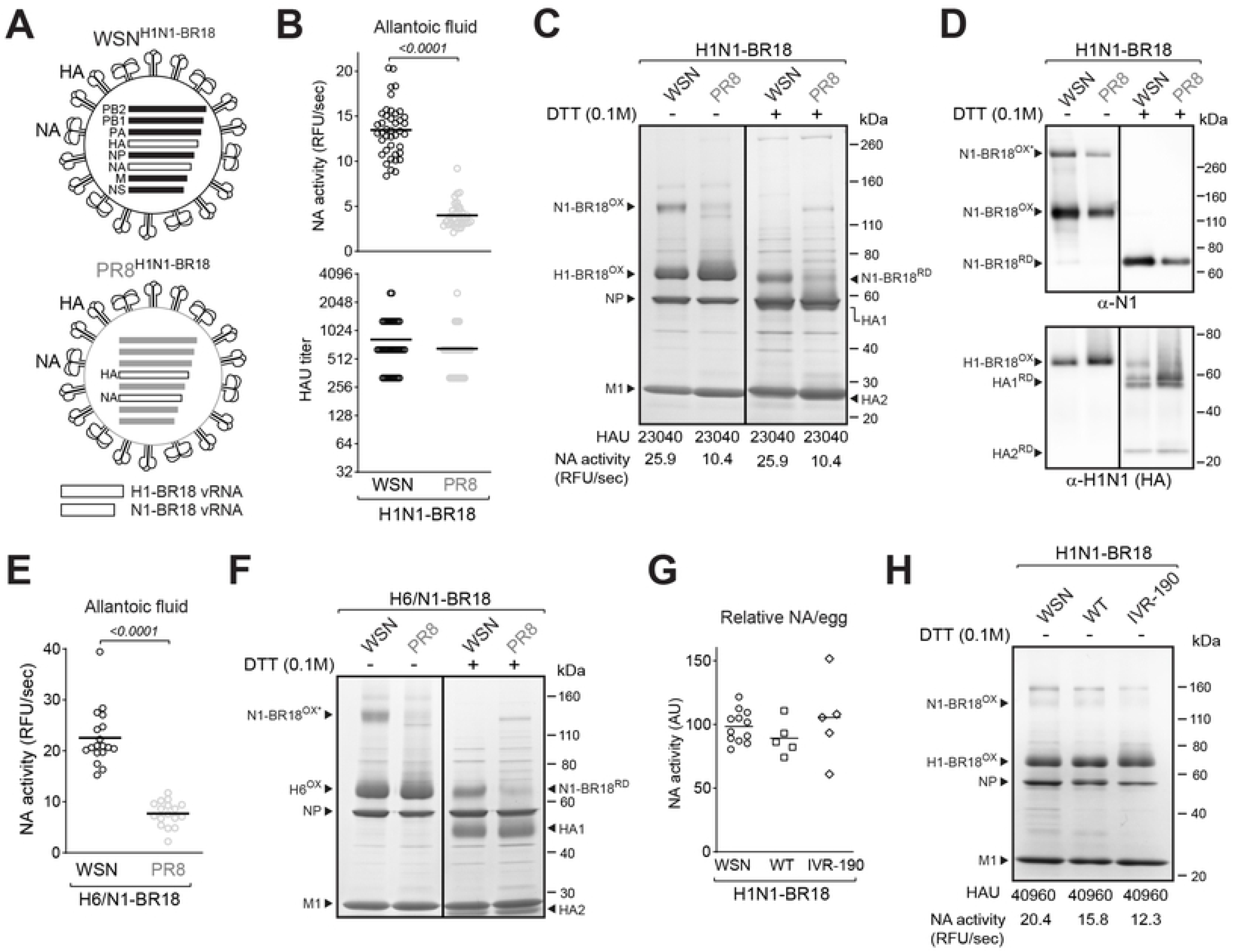
The WSN backbone supports high N1 virion incorporation independent of HA. **A.** Diagram of the double-gene reassortant IAVs with WSN and PR8 backbones and the HA and NA gene segments from the H1N1 strain A/Brisbane/02/2018. **B.** Eggs were harvested 3 days post-infection and the HAU titers and NA activities were measured in each egg using equal allantoic fluid volumes. Individual egg data from three independent experiments with different batches of eggs are shown with the mean (bar). Significant *P* values calculated from a two-tailed unpaired t-test (95% CI) are displayed. **C.** Coomassie stained gel of the purified BPL inactivated virions (5 μg per lane), treated with DTT as indicated. Oxidized and reduced N1-BR18 and H1-BR18 are indicated with NP and M1. HAU titres and NA activity values are means from the 3 independently purified virus batches with equal total protein concentrations (1 mg/ml). **D.** NA (top) and HA (bottom) immunoblots of the BPL inactivated purified virions (0.5 μg per lane), untreated or reduced with DTT prior to SDS-PAGE. Bands corresponding to oxidized and reduced N1-BR18 and H1-BR18 are indicated. Asterisk denotes SDS resistant NA homo-tetramers. Note the H1N1 antisera shows some NA reactivity. **E.** Eggs were harvested 3 days post-infection and the NA activities were measured in each egg using equal allantoic fluid volumes. Individual egg data from two independent experiments with different batches of eggs are shown with the mean (bar) and the *P* value from a two-tailed unpaired t-test (95% CI). **F.** Coomassie stained gel of the purified BPL inactivated virions (5 μg per lane) treated with DTT as indicated. Oxidized and reduced forms of N1-BR18 and H6 are indicated with NP and M1. **G.** Eggs were harvested 3 days post-infection with WSN^H1N1-BR18^, the H1N1 strain isolate A/Brisbane/02/2018 (WT) and CVV (IVR-190). The relative total amount of NA from each egg is shown with the mean (bar). **H.** Coomassie stained gel showing the purified BPL inactivated virions (5 μg per lane). Oxidized and reduced forms of N1-BR18 and H1-BR18 are indicated with NP and M1. HAU titres and NA activity values from the purified virus batches with equal total protein concentrations (1 mg/ml) are displayed.

### Changing the CVV backbone can provide advantages for NA virion content

Most seasonal viral-based influenza vaccines are produced in eggs using either the WHO recommended wild type field strain or a CVV that has been generated with higher growth properties [14]. Therefore, we benchmarked the NA levels produced by the WSN^H1N1-BR18^ virus against the wild-type (WT) H1N1 BR-18 field isolate as well as the BR18 CVV (IVR-190) approved for the manufacturing of influenza vaccines for the 2019-2020 season. Based on the relative NA activity per egg the WSN^H1N1-BR18^ virus generated NA levels that were on par or better than WT and slightly lower than IVR-190 (Fig. 4G). Following isolation and normalization for total protein levels, the NA content and activity in WSN^H1N1-BR18^ virions was found to be slightly higher than the WT strain and almost twice as high as IVR-190 (Fig. 4H), however the IVR190 strain did produce higher viral yields. This data further supports that influenza viruses can accommodate a range of NA and HA ratios for a given pair and suggests that the NA content in CVVs used to produce influenza vaccines can be increased.

### Plasticity in the HA and NA balance allows for increase in NA virion incorporation

Changes in the virion NA to HA ratio suggests that the functional balance between a particular NA and HA pair has some level of plasticity. To investigate the functional balance, we examined the WSN^H1N1-BR18^ and PR8^H1N1-BR18^ viruses using the NA inhibitor zanamivir in conjunction with a bio-layer interferometer (BLI) equipped with a streptavidin biosensor containing a biotinylated 2,3-sialic acid analogue receptor, Neu5Ac-α2-3Gal-β1-4GlcNAc-β-PAA-biotin (3’SLN). Initially, we confirmed that the zanamivir 50% inhibitory concentration (IC_50_) for the purified BPL inactivated viruses were similar (Fig. 5A) and that the IC_50_ values were in line with previous studies using H1N1 IAVs [37]. When the 3’SLN association was monitored by BLI, the HA-receptor binding of both viruses was disrupted by NA within a few seconds and the disruption could be overcome by zanamivir in a dose-dependent manner (Fig. 5B). However, higher zanamivir concentrations were required for efficient 3’SLN binding of the WSN^H1N1-BR18^ virus (Fig. 5B), presumably because the virus possesses more NA, requiring higher zanamivir concentration for complete NA activity inhibition (Fig. 5A, inset).

**Figure 5.**
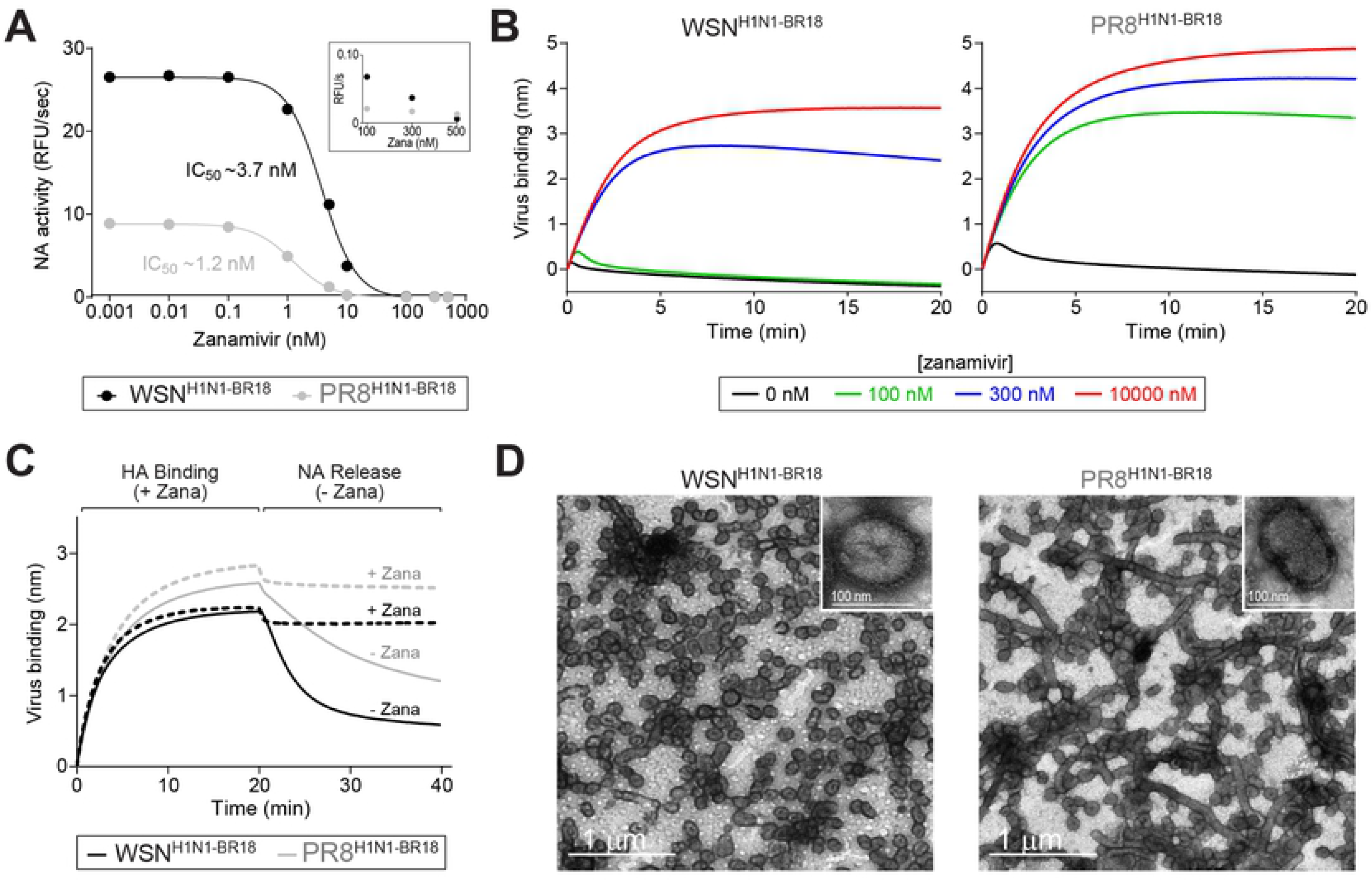
Viral properties that differ with changes in the NA to HA ratio. **A.** NA activity in the purified BPL inactivated viruses were measured using MUNANA in the presence of increasing zanamivir concentrations to calculate the 50% inhibitory concentration (IC_50_). Measurements were made using 0.5 μg total viral protein. Inset shows the NA activity values at high zanamivir concentrations. **B.** Binding of purified WSN^H1N1-BR18^ virions (left graph) and PR8^H1N1-BR18^ virions (right graph) to 3’-SLN were measured in the presence of the indicated zanamivir concentrations by biolayer interferometry (BLI). Each binding curve was generated with a biosensor containing the same density of immobilized 3’SLN and 0.1 mg/ml of the purified viruses. **C.** HA binding of the purified virions (0.1mg/ml) to 3’SLN and NA-mediated release of the virus bound to 3’-SLN were measured by BLI. HA-binding was monitored for 20 min in the presence of 10 μM zanamivir to inhibit NA activity. The biosensor containing the bound virus was moved to a new well with (dashed line) or without zanamivir (line) and the NA-mediated release was followed. **D.** Negative stained transmission electron microscopy images of BPL inactivated purified WSN^H1N1-BR18^ and PR8^H1N1-BR18^ virions. Insets contain higher magnification images of a spherical virion. Scale bars (white) are included.

The BLI results indicate that either the low HA content and/or higher NA content altered the HA-mediated binding of WSN^H1N1-BR18^ compared to PR8^H1N1-BR18^. To separate these two processes, HA-mediated binding to 3’SLN was monitored in the presence of high zanamivir concentrations and NA-mediated release was examined by removing the inhibitor. In the presence of zanamivir, both viruses showed similar HA binding with the WSN^H1N1-BR18^ virus displaying lower saturation levels (Fig. 5C). In contrast, the NA mediated release of the WSN^H1N1-BR18^ virus was substantially faster than the PR8^H1N1-BR18^ virus (Fig. 5C), confirming that the NA virion increase is reflected in a functional analysis. These results indicate that IAVs can support variations in the HA and NA ratio of a given pair that result in different functional balances.

### Viruses with the WSN backbone and higher NA content are predominantly spherical

NA has been proposed to cluster on the viral surface at regions of high membrane curvature [6, 38, 39]. Therefore, we examined the morphology of the purified BPL inactivated WSN^H1N1-BR18^ and PR8^H1N1-BR18^ virions by electron microscopy. The WSN^H1N1-BR18^ virions displayed a homogeneous spherical morphology with a diameter of ~100 nm, whereas the PR8^H1N1-BR18^ virions showed a mixture of spherical and filamentous morphologies, with the later having lengths greater than 1 μm (Fig. 5D). While it is interesting to speculate that membrane crowding causes or results in the higher NA content within the spherical virions, it is equally plausible that the morphology difference is an inherent property of the WSN backbone or a partial contributing factor to the higher NA content in the virions.

### Increasing the NA virion content can balance the NA and HA antibody response

To determine if the increase in NA virion content we achieved can improve the NA antibody response, a comparative immunization analysis was performed using mice (Fig. 6A). Prior to the immunization experiment we used Coomassie stained gel densitometry to estimate the amount of NA in the inactivated purified WSN^H1N1-BR18^ and PR8^H1N1-BR18^ virions (Fig. 6B). Based on the standard curve from recombinant N1-BR18 protein, 5 μg of total WSN^H1N1-BR18^ and PR8^H1N1-BR18^ viral protein were estimated to contain ~750 ng and ~225 ng of N1-BR18, respectively, a ratio in-line with the NA activity difference (Figs. 6B and 4C). Mice were then intramuscularly immunized with 1 or 5 μg of purified inactivated WSN^H1N1-BR18^ or PR8^H1N1-BR18^ virions in parallel. One cohort received one dose while the other received a second dose after 21 days and sera were collected 21 days following the last immunization.

**Figure 6.**
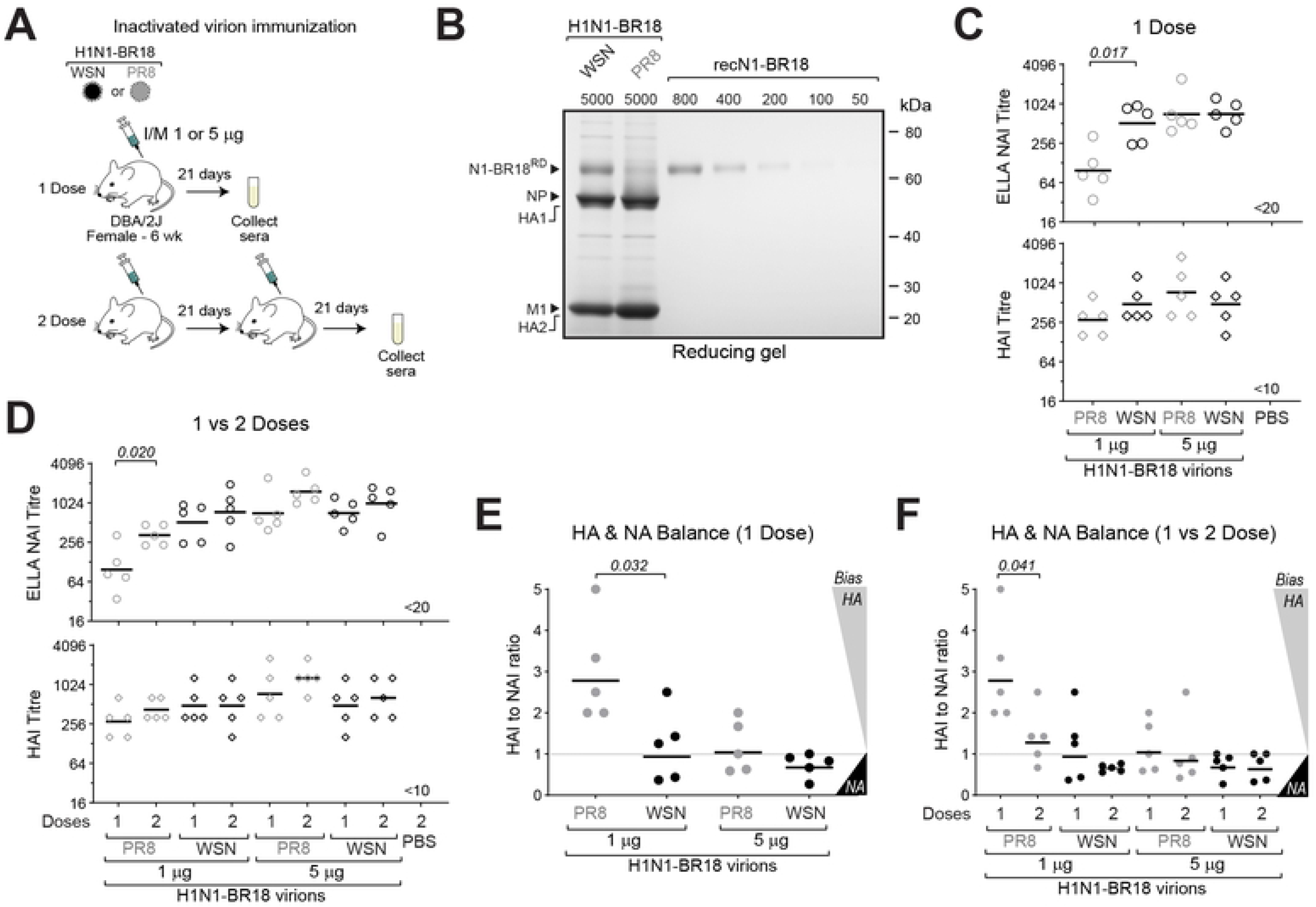
Virions with increased NA content elicit a more balanced NA and HA response. **A.** Reducing Coomassie stained gel that was used to estimate the NA protein content in the BPL inactivated purified WSN^H1N1-BR18^ and PR8^H1N1-BR18^ virions by densitometry. The indicated amounts of recombinant N1-BR18 were used to generate the standard curve. **B.** Diagram depicting the one and two dose immunization schemes in mice using two different amounts (1 μg and 5 μg) of BPL inactivated purified WSN^H1N1-BR18^ and PR8^H1N1-BR18^ virions. **C.** Graphs displaying the ELLA NAI titres (Top) and the HAI titres (Bottom) obtained from mice following one dose with the indicated virion amounts. Only significant *P* values comparing the different virions at each dose are displayed. **D.**Graphs comparing the ELLA NAI titres (Top) and HAI titres (Bottom) in the mice that received one and two doses of the indicated virion amounts. Only significant *P* values comparing one and two doses for each virion and protein amount are displayed. **E.** The HAI to NAI titre ratio was plotted for each serum from the mice that received one dose of the indicated amounts of the virions. Values >>1 indicate the antibody response is HA biased, values ~1 indicate a more balanced response and values <<1 indicate a NA biased response. Only significant *P* values comparing the different virions at each dose are displayed. **F.** Graph comparing the HAI to NAI titre ratio for each serum from the one and two dose immunizations for each virion and protein amount. Only significant *P* values comparing one and two doses for each virion and amount are displayed.

The enzyme-linked lectin assay (ELLA) revealed that the NA inhibitory (NAI) titres from mice following a single 1 μg dose of WSN^H1N1-BR18^ virions were ~5 fold higher than those from 1 μg of PR8^H1N1-BR18^ virions and were on par with the 5 μg dose of both virions, whereas the hemagglutination inhibition (HAI) titres were similar across the groups (Fig. 6C). The second dose only showed improvements in NAI titres for the 1 μg dose of PR8^H1N1-BR18^ virions, which contained the least amount of NA protein, and the HAI titres remained largely unchanged by the second dose (Fig. 6D). When we determined the HAI to NAI ratio in each mouse, a clear trend towards a more balanced response was observed with increasing NA virion content in the single dose immunizations as the ratio coalesced towards 1 (Fig. 6E). This observation was further supported in the mice that received the second 1 μg dose of PR8^H1N1-BR18^ virions, which caused the ratio to decrease from ~2.5 (HA biased) to ~ 1.25 (more balanced), and the second 1 μg dose of WSN^H1N1-BR18^ virions, which resulted in all the ratios coalescing to slightly below 1 (Fig. 6F). Together, these results imply that attainable increases in NA virion content can improve the balance between the NA and HA inhibitory antibody responses, especially at lower antigen concentrations, indicating this potentially beneficial response can be achieved without dramatically increasing the HA content in a viral-based vaccine dose.

## Discussion

Influenza NA is a labile, homotetrameric Ca^2+^-dependent enzyme with a complex maturation process that is coordinated by the N-terminal transmembrane region and the C-terminal enzymatic head domain [28, 36, 40–43]. Recently, it was shown that each NA monomer in a tetramer functions independently (noncooperatively) and that the low stability of the Ca^2+^-dependent oligomeric conformation varies between strains of the same subtype [29]. These nonideal antigen properties and the limited abundance, likely impact both the quantity and quality of the NA in viral-based influenza vaccines, hindering a productive antibody response [15–17]. As influenza vaccine efficacy remains less than optimal, there are significant efforts to enhance the NA response to create vaccines that provide higher efficacy and broader cross protection against strains [44].

Our results demonstrate that the bottleneck of NA abundance in H1N1 virions can be addressed by changing the internal gene backbone used to generate CVVs. This approach preserves the NA and HA antigens and the increase in NA content was shown to induce higher NAI titres without affecting the HAI titres, resulting in a more balanced NA and HA antibody responses. The improved balance against both antigens was more evident in the single immunizations with the lowest virion amounts, suggesting the NA antibody response in mice is limited by a quantity threshold rather than HA immunodominance. Future challenge experiments using immunizations with even lower virion amounts will help to determine if the increase in NA content can decrease the antigen amount required for protection, and to define the NA antigen amounts in mice that correlate with protection from strains with unmatched HAs.

Although WSN is a commonly used highly adapted lab strain like PR8, it has not been explored much as a CVV backbone because of many reports showing it is neurovirulent in mice [45–47]. However, the neurovirulence trait has been linked to the NA in WSN [48, 49], requires intracerebral inoculation [45], and in CVVs both the HA and NA gene segments are replaced during generation. In addition, we have observed no clinical symptoms (*e.g.* fever, weight loss, nasal symptoms or low energy) in ferrets that were inoculated with a series of identical HA and NA double-gene WSN and PR8 reassortants for another study. We are not certain that using classical reassortment to create a CVV with a WSN backbone will improve viral growth and retain the higher NA virion content, but our results showing that the NA virion content is higher than an approved CVV and the observation that viral protein yields are on par with the WT field isolate, which is used to benchmark related CVVs, suggest further investigation is warranted.

The neurovirulence requirement for the NA from WSN has been attributed to its unique ability to enable trypsin-independent viral replication by facilitating the recruitment of the protease plasminogen for HA processing [50, 51]. Although not required, the M and NS gene segments from WSN were also shown to promote viral growth in neurons [48, 49], which possess especially high sialic acid content [52], suggesting these segments could potentially contribute to the increased NA virion content. In line with this possibility, the virions with the WSN backbone displayed a more homogeneous spherical morphology, a phenotype that has previously been associated with the M gene segment [53–55]. Mechanistically, spheres could potentially accommodate more copies of asymmetrical shaped NA tetramers which have been proposed to localize to membrane regions with high curvature [38, 39, 55]. A second difference was observed during purification where the WSN virion pellet showed a propensity to flocculate and foam during resuspension, indicating there potentially is lipid differences between the virions. However, the cause and effect remain unclear because it could result from an internal gene product changing the cell host membrane composition or from the increased NA content leading to the recruitment of specific lipids. By identifying the internal gene segment or constellation that promotes NA virion incorporation it should be possible to discern if any of these properties are the main contributing factors and if other seasonal IAV gene segments promote even higher NA incorporation.

While our data indicate that virions can support a range of functional balances for the same NA and HA, raising the question of whether changes in the balance can contribute to influenza pathogenicity or transmission *in vivo*. Despite this flexibility in the functional balance, there is likely an upper limit in the NA content a virion can possess and the HA influence on the NA content in a CVV for a vaccine is unavoidable. The associated decrease in the virion HA content upon the increase in NA, which is an expected outcome due to mass action and the packaging limitations, is also not ideal for maximizing the vaccine dose number unless the added benefit from NA can improve the efficacy and/or decrease the required dose amount. These critical questions emphasize the need for a systematic analysis of the requirements for protection by NA and HA independently, and together, to determine if the theoretical upper NA limit in a virion is enough to overcome the NA quantity bottleneck in current influenza vaccines that are dosed based on HA.

## Materials and Methods

### Reagents and antibodies

Dulbecco’s Modified Eagles Medium (DMEM), fetal bovine serum (FBS), L-glutamine, penicillin/streptomycin (P/S), Opti-MEM I (OMEM), anti-goat IgG HRP-linked secondary antibody, Simple Blue Stain, Novex 4-12% Tris-Glycine SDS-PAGE gels, dithiothreitol (DTT) and Lipofectamine 2000 transfection reagent were all purchased from Thermo Fisher Scientific. Zanamivir and 2’-(4-methylumbelliferyl)-α-d-*N*-acetylneuraminic acid (MUNANA) were acquired from Moravek Inc and Cayman Chemicals, respectively. β-propiolactone and formaldehyde were purchased from Sigma. Anti-rabbit IgG HRP-linked secondary antibody and 0.45-μm polyvinylidene difluoride (PVDF) membrane were obtained from GE healthcare. Specific-Pathogen-Free (SPF) eggs and turkey red blood cells (TRBCs) were purchased from Charles River Labs and the Poultry Diagnostic and Research Center (Athens, GA), respectively. The H1N1 A/Brisbane/02/2018 field isolate (WT) and CVV (IVR-190) were kindly provided by the WHO. Rabbit Antisera against NA was generated by Agrisera (Sweden) using NA-WSN residues 35–453 isolated from *E. coli* inclusion bodies [29]. Polyclonal goat antiserum against the H1N1 influenza virus A/Fort Monmouth/1/1947 (NR-3117) was obtained from BEI Resources, NIAID, NIH.

### Plasmids and constructs

The eight reverse genetics (RG) plasmids encoding the WSN33 and PR8 gene segments were provided by Dr. Robert Webster (St. Jude Children’s Research Hospital). The RG plasmids were sequenced before use and correspond with the following GenBank Identifications: LC333182.1 (WSN33-PB2), LC333183.1 (WSN33-PB1), LC333184.1 (WSN33-PA), LC333185.1 (WSN33-HA), LC333186.1 (WSN33-NP), LC333187.1 (WSN33-NA), MF039638.1 (WSN33-M) LC333189.1 (WSN33-NS), CY038902.1 (PR8-PB2), CY038901.1 (PR8-PB1), CY084019.1 (PR8-PA), CY146825.1 (PR8-HA), CY038898.1 (PR8-NP), CY038897.1 (PR8-NA), MH085246.1 (PR8-M), and CY038899.1 (PR8-NS). The RG plasmids containing the NA (N1-CA09) and the H6 gene (A/turkey/Mass/3740/1965) were described previously [28, 56]. To generate the NA (N1-BR18) and HA (H1-BR18) RG plasmids, the NA and HA gene segments from IVR-190, grown in SPF chicken eggs, were amplified by RT-PCR and cloned into the pHW2000 plasmid using restriction enzymes [57]. All constructs were confirmed by sequencing (FDA core facility or Macrogen).

### Cell culture and viral reverse genetics

Madin-Darby canine kidney 2 (MDCK.2; CRL-2936) cells and HEK 293T/17 cells (CRL-11268) were both obtained from LGC Standards and cultured at 37 °C with 5% CO_2_ and ~95% humidity in DMEM containing 10% FBS and 100 U/ml P/S. Reassortant viruses were created by 8-plasmid reverse genetics [57] in 6-well plates using the indicated NA, or NA and HA pair, and the complimentary seven, or six, ‘backbone’ gene segments of WSN or PR8. For each virus, ~1 × 10^6^ 293T and ~1 x 10^6^ MDCK.2 cells were plated in a well the day before. The next day, the eight plasmids (1 μg of each) were added to 200 μl of OMEM, mixed with 18 μl of lipofectamine, and incubated 45 min at room temperature. After the incubation, the cells were washed with 2 ml OMEM, the mixture was added to the well and incubated 5 min at 37 °C before receiving 800 μl OMEM. At ~24 h post-transfection, 1 ml OMEM containing 4 μg/ml TPCK trypsin was added to each well. Rescued viruses in the culture medium were harvested ~96 h post-transfection, clarified by sedimentation (2000 × g; 5 min) and passaged using SPF eggs or MDCK.2 cells.

### Viral passaging and analysis in MDCK cells

Reverse genetics rescued viruses in culture medium were diluted 1:1,000 in infection media (DMEM with 0.1% FBS, 0.3% BSA, 1% P/S and 2 μg/ml of TPCK trypsin) and 10 ml was added to a T75cm flask containing MDCK.2 cells at ~95% confluency. The cells were incubated 3 days at 37 °C prior to harvesting. The virus-containing culture medium was clarified by centrifugation (2,000 × g; 5 min), aliquoted, stored at −80 °C, and the median tissue culture infectious dose (TCID_50_ ml^−1^) was determined by cytopathic effects (CPE) in MDCK.2 cells at 72 h [58]. For the NA expression analysis, T25cm flasks with ~95% confluent MDCK.2 cells were infected with the viruses at an MOI ~0.001 and processed as follows. Cells were incubated with virus diluted in infection media for 30 min at 4 °C. Unbound virus was removed and 5 ml of fresh infection media was added. Infections were monitored by observing CPE and the viruses and cells were harvested by scraping once the infection completed, ~3 days. The cell remains were isolated by sedimentation (5 min at 10,000 x g), virus containing media was transferred to a new tube and the cell pellet was lysed in 500 μl of lysis buffer (20 mM Tris, pH 7.4, 150 mM NaCl, 1% *n*-dodecyl *β*-D-maltoside, 1X protease inhibitor (Sigma)). Lysates were sonicated on ice 30 s, sedimented (20,000 × *g*, 5 min), and the post-nuclear supernatants were retained. NA activity was measured using 50 μl of medium and 5 μl of the post nuclear supernatant, whereas HAU titres were determined using 50 μl of medium.

### Viral passaging in SPF chicken eggs

Initial passages (E1) were carried out by inoculating 9-11 day old embryonic SPF chicken eggs with 100 μl of the rescued virus in culture medium. Eggs were incubated for 3 days at 33 °C and placed at 4 °C for 2 h prior to harvesting the allantoic fluid. The allantoic fluid was clarified by centrifugation (2000 × g; 5 min), aliquoted and stored at −80 °C. The second passage was carried out similarly. E1 aliquots of the viruses being compared were thawed at 37 °C, diluted 1:1000 in sterile PBS and 100 μl was used to inoculate groups of eight or more 9-11 day old embryonic eggs. The allantoic fluid was harvested individually from each egg at 3 days post-infection, clarified by centrifugation (2000 × g; 5 min), and NA activity and HAU measurements were taken prior to inactivation and purification of the virions.

### Viral inactivation by β-propiolactone and formaldehyde

To analyse potential affects on NA and HA, 1 ml aliquots of the WSN^N1-BR18^ and PR8^N1-BR18^ viruses in allantoic fluid were treated with the indicated concentrations of β-propiolactone (BPL) or formaldehyde (FA) for ~16 h at 4 °C. Following the treatment, NA activity was measured in each sample using 10 μl of allantoic fluid and HAU titres were measured using 50 μl of allantoic fluid. For inactivation prior to purification, the viruses in allantoic fluid were treated with 0.05% BPL for ~16 h at 4 °C.

### Viral purification

BPL treated viruses in allantoic fluid were first isolated by sedimentation (100,000 × g; 45 min) at 4 °C through a sucrose cushion (25% w/v sucrose, PBS pH 7.2 and 1 mM CaCl_2_) equal to 12.5% of the sample volume. The pelleted virions were resuspended in 500 μl sucrose 12.5% w/v in PBS pH 7.2 containing 1 mM CaCl_2_, layered on top of a discontinuous sucrose gradient containing four 8.5 ml sucrose layers (60% w/v, 45% w/v, 30% w/v and 15% w/v in PBS pH 7.2 and 1 mM CaCl2) and centrifuged at 100,000 × g for 2 h at 4 °C. Fractions were collected and the density was determined using a refractometer. Fractions with a density between 30-50% sucrose w/v were pooled, mixed with 2 volumes of PBS pH 7.2 and 1 mM CaCl_2_, and sedimented (100,000 × g; 45 min). The supernatant was discarded, the sedimented virions were resuspended in 250 μl PBS pH 7.2 containing 1 mM CaCl_2_ and the total protein concentration was determined using a BCA protein assay kit (Pierce) according to the 96-well plate protocol. All purified viruses were adjusted to a concentration of 1 mg/ml using PBS pH 7.2 containing 1 mM CaCl_2_, NA activity was measured using 0.5 μg and HAU titres were determined.

### NA activity measurements, IC_50_ determination and HAU titres

All NA activity measurements were performed in a 96-well low protein binding black clear bottom plate (Corning). Each sample was mixed with 37°C reaction buffer (0.1 M KH_2_PO_4_ pH 6.0 and 1 mM CaCl_2_) to a volume of 195 μl. Reactions were initiated by adding 5 μl of 2 mM MUNANA and the fluorescence was measured on Cytation 5 (Biotek) plate reader at 37°C for 10 min using 30 sec intervals and a 365 nm excitation wavelength and a 450 nm emission wavelength. Final activities were determined based on the slopes of the early linear region of the emission versus time graph. The zanamivir IC_50_ concentrations were determined similarly by including the indicated zanamivir concentrations in the reaction using 0.5 μg of purified viral protein. The slopes were then plotted against each zanamivir concentration and the IC_50_ value was determined using a variable slope four parameter nonlinear fit analysis in Graphpad Prism 8. HAU titres were determined by a two-fold serial dilution in a 96-well plate using a sample volume of 50 μl. Following the dilution, 50 μl of 0.5% TRBCs were added to each well and the plate was incubated 30 min at room temperature. HAU titres were determined as the last well where agglutination was observed.

### SDS-PAGE, Coomassie staining, immunoblotting, and densitometry

Purified virions equalling 5 μg of total viral protein (Coomassie), or 100 ng (immunoblots), were mixed with 2X sample buffer containing 0.1 M DTT as indicated. Samples were heated at 50 °C for 5 min and resolved using a 4-12 % polyacrylamide Tris-Glycine SDS-PAGE wedge gel. Gels were either stained with simple blue or transferred to a 0.45-μm pore PVDF membrane at 65 V for 1 h. PVDF membranes were blocked with milk/PBST (3% nonfat dry milk, PBS, pH 7.4, 0.1% Tween 20) for 30 min and processed with the indicated antibodies and appropriate HRP-linked secondary antibody. Immunoblots were developed with the SuperSignal West Femto kit (ThermoFischer) and imaged using an Azure C600 or a Syngene G Box. The same systems were used for Coomassie gel imaging. NA protein amounts in 5 μg of purified virions was determined by Coomassie gel densitometry analysis using purified recombinant N1-BR18 as a standard. Samples were resolved on the same 4-12% reducing SDS-PAGE gel, stained with simple blue, and imaged with an Azure C600. Densities of the NA bands were analysed using ImageJ and the protein amount was calculated using the standard curve from the recombinant protein samples.

### Bio-Layer Interferometry

Binding of the purified BPL inactivated virions to the 2,3-sialic acid analogue receptor, Neu5Ac-α2-3Gal-β1-4GlcNAc-β-PAA-biotin (3’SLN) was performed at 25℃ using a ForteBio Octet Red96 bio-layer interferometer equipped with ForteBio Streptavidin biosensors at 25 °C and a plate shake speed of 1000 rpm. Baselines were established for 2 min in 200 μl of buffer (PBS pH 7.4 containing 1mM CaCl_2_) prior to and post incubation of the biosensor with 200 μl of 6.7 nM 3’SLN for 10 min. Biosensors were then transferred to wells containing 200 μl of 0.1 mg/ml WSN^H1N1-BR18^ or PR8^H1N1-BR18^ virions in buffer with the indicated concentrations of zanamivir for 20 min. HA-mediated association was analysed by incubating the 3’SLN loaded biosensors in 200 μl of 0.1 mg/ml virions and 10 μM zanamivir for 20 min. NA-mediate dissociation was examined by transferring the biosensors to new wells that contained 200 μl of buffer for 20 min. Transfer to a well containing buffer with 10 μM zanamivir was included for a NA-inhibited dissociation control.

### EM analysis

Purified BPL inactivated virions in PBS pH 7.4, 1 mM CaCl_2_ with a total viral protein concentration of 1 mg/ml were diluted 1:4 in PBS pH 7.4 and 15 μl were spotted on formvar/carbon film 300 copper grids. Following a 5 min incubation, the virions were fixed with 2.5% Glutaraldehyde in 0.1 M PBS buffer pH 7.3 for 3 min, washed 3 times with dH_2_O, and stained with 2% Phosphotungstic Acid (PTA) for 50 sec. For each step the solution was drawn off the edge of the grid with filter paper. Grids were placed directly into a grid box and air-dried overnight before imaging with a Jeol 1400 transmission electron microscope (TEM) equipped with a Gatan US 1000XP digital camera.

### Ethics statement

All animal experiments were approved by the U.S. FDA Institutional Animal Care and Use Committee (IACUC) under Protocol #2003–18. The animal care and use protocol meets National Institutes of Health (NIH) guidelines.

### Immunizations, HAI and NAI titre measurements

Female DBA/2J mice were purchased from The Jackson Labs (#800671). At ~6 weeks of age, mice were immunized intramuscularly with the indicated protein amounts of purified BPL inactivated virions in 50 μl PBS. Mice receiving a second dose were immunized similarly 21 days after the first dose and blood was collected 21 days following the last immunization in Vacutainer SST II Advance Tubes (BD). The tubes were incubated at room temperature for ~1 h and placed at 4 °C overnight. The next day the tubes were centrifuged (5 min; 2,000 × g) and the sera was transferred to a new tube and stored at −80 °C for further analyses. A portion of the sera was treated with receptor destroying enzyme (RDE) overnight at 37 °C and then heat inactivated at 56 °C for 45 min. The HAI assay was performed in a 96-well plate using the PR8^H1N1-BR18^ virus in allantoic fluid. The virus was diluted to a titre of 4 HAUs and incubated with a two-fold serial dilution of the RDE treated sera in a total volume of 50 μl for 30 min at room temperature. Each well then received 50 μl of 0.5% TRBCs in PBS pH 7.2 and the plate was incubated for 45 min at room temperature. The HAI titre was calculated based on the sera dilution corresponding to the last well that prevented hemagglutination. NAI titres were determined using the enzyme-linked lectin assay (ELLA) with bovine fetuin and the PR8^H6/N1-BR18^ virus in allantoic fluid as described previously [59, 60].

### Statistical analysis

Data analysis was performed using GraphPad Prism 8 software. The P values were calculated using Student’s unpaired t-test with a two-tailed analysis and a confidence interval (CI) of 95% under the assumptions that the data from the independent biological replicates follow a Gaussian distribution with equal standard deviation between sample groups.

## Acknowledgements

We would like to thank members of the Division of Viral Products at the Center for Biologics Evaluation and Research for offering several helpful suggestions and insightful discussions. This work was supported in part by federal funds from the NIAID, National Institutes of Health, Department of Health and Human Services, under CEIRS contract number HHSN272201400005C.

## Author contributions

J.G. performed most of the experiments and analysed the data. H.W. designed and performed the immunization experiments with J.G. X.L. designed, performed, and interpreted the BLI data. M.R.M. analysed the virions with H6. Y.G. processed and imaged the virions by TEM. R.D. conceived and supervised the study with help from Z.Y., and all authors contributed to the writing of the manuscript.

## References

1. Zhao Q, Potter CS, Carragher B, Lander G, Sworen J, Towne V, et al. Characterization of virus-like particles in GARDASIL(R) by cryo transmission electron microscopy. Hum Vaccin Immunother. 2014:10(3):734–9. Epub 2013/12/05. doi: 10.4161/hv.27316. PubMed PMID: 24299977: PubMed Central PMCID: PMCPMC4130261.

2. Murphy BR, Kasel JA, Chanock RM. Association of serum anti-neuraminidase antibody with resistance to influenza in man. N Engl J Med. 1972:286(25):1329–32. Epub 1972/06/22. doi: 10.1056/NEJM197206222862502. PubMed PMID: 5027388.

3. Couch RB, Kasel JA, Gerin JL, Schulman JL, Kilbourne ED. Induction of partial immunity to influenza by a neuraminidase-specific vaccine. J Infect Dis. 1974:129(4):411–20. PubMed PMID: 4593871.

4. Johansson BE, Bucher DJ, Kilbourne ED. Purified influenza virus hemagglutinin and neuraminidase are equivalent in stimulation of antibody response but induce contrasting types of immunity to infection. J Virol. 1989:63(3):1239–46. Epub 1989/03/01. PubMed PMID: 2915381: PubMed Central PMCID: PMC247820.

5. Treanor JJ, Schiff GM, Hayden FG, Brady RC, Hay CM, Meyer AL, et al. Safety and immunogenicity of a baculovirus-expressed hemagglutinin influenza vaccine: a randomized controlled trial. JAMA. 2007:297(14):1577–82. Epub 2007/04/12. doi: 10.1001/jama.297.14.1577. PubMed PMID: 17426277.

6. Harris A, Cardone G, Winkler DC, Heymann JB, Brecher M, White JM, et al. Influenza virus pleiomorphy characterized by cryoelectron tomography. Proc Natl Acad Sci U S A. 2006:103(50):19123–7. doi: 10.1073/pnas.0607614103. PubMed PMID: 17146053: PubMed Central PMCID: PMCPMC1748186.

7. Zheng A, Sun W, Xiong X, Freyn AW, Peukes J, Strohmeier S, et al. Enhancing Neuraminidase Immunogenicity of Influenza A Viruses by Rewiring RNA Packaging Signals. J Virol. 2020:94(16). Epub 2020/06/05. doi: 10.1128/JVI.00742-20. PubMed PMID: 32493826: PubMed Central PMCID: PMCPMC7394900.

8. Mitnaul LJ, Castrucci MR, Murti KG, Kawaoka Y. The cytoplasmic tail of influenza A virus neuraminidase (NA) affects NA incorporation into virions, virion morphology, and virulence in mice but is not essential for virus replication. J Virol. 1996:70(2):873–9. Epub 1996/02/01. PubMed PMID: 8551626: PubMed Central PMCID: PMC189890.

9. Mitnaul LJ, Matrosovich MN, Castrucci MR, Tuzikov AB, Bovin NV, Kobasa D, et al. Balanced hemagglutinin and neuraminidase activities are critical for efficient replication of influenza A virus. J Virol. 2000:74(13):6015–20. Epub 2000/06/14. PubMed PMID: 10846083: PubMed Central PMCID: PMC112098.

10. Kaverin NV, Matrosovich MN, Gambaryan AS, Rudneva IA, Shilov AA, Varich NL, et al. Intergenic HA-NA interactions in influenza A virus: postreassortment substitutions of charged amino acid in the hemagglutinin of different subtypes. Virus Res. 2000:66(2):123–9. Epub 2000/03/22. doi: 10.1016/s0168-1702(99)00131-8. PubMed PMID: 10725545.

11. Kaverin NV, Gambaryan AS, Bovin NV, Rudneva IA, Shilov AA, Khodova OM, et al. Postreassortment changes in influenza A virus hemagglutinin restoring HA-NA functional match. Virology. 1998:244(2):315–21. Epub 1998/05/28. doi: 10.1006/viro.1998.9119. PubMed PMID: 9601502.

12. Wagner R, Matrosovich M, Klenk HD. Functional balance between haemagglutinin and neuraminidase in influenza virus infections. Rev Med Virol. 2002:12(3):159–66. doi: 10.1002/rmv.352. PubMed PMID: 11987141.

13. Houser K, Subbarao K. Influenza vaccines: challenges and solutions. Cell host & microbe. 2015:17(3):295–300. doi: 10.1016/j.chom.2015.02.012. PubMed PMID: 25766291: PubMed Central PMCID: PMCPMC4362519.

14. Gerdil C. The annual production cycle for influenza vaccine. Vaccine. 2003:21(16):1776–9. Epub 2003/04/11. doi: 10.1016/s0264-410x(03)00071-9. PubMed PMID: 12686093.

15. Johansson BE, Matthews JT, Kilbourne ED. Supplementation of conventional influenza A vaccine with purified viral neuraminidase results in a balanced and broadened immune response. Vaccine. 1998:16(9-10):1009–15. PubMed PMID: 9682352.

16. Wohlbold TJ, Nachbagauer R, Xu H, Tan GS, Hirsh A, Brokstad KA, et al. Vaccination with adjuvanted recombinant neuraminidase induces broad heterologous, but not heterosubtypic, cross-protection against influenza virus infection in mice. MBio. 2015:6(2):e02556. doi: 10.1128/mBio.02556-14. PubMed PMID: 25759506: PubMed Central PMCID: PMCPMC4453582.

17. Chen YQ, Wohlbold TJ, Zheng NY, Huang M, Huang Y, Neu KE, et al. Influenza Infection in Humans Induces Broadly Cross-Reactive and Protective Neuraminidase-Reactive Antibodies. Cell. 2018:173(2):417–29 e10. doi: 10.1016/j.cell.2018.03.030. PubMed PMID: 29625056: PubMed Central PMCID: PMCPMC5890936.

18. Dou D, Revol R, Östbye H, Wang H, Daniels R. Influenza A Virus Cell Entry, Replication, Virion Assembly and Movement. Frontiers in Immunology. 2018:9(1581). doi: 10.3389/fimmu.2018.01581.

19. Ostbye H, Gao J, Martinez MR, Wang H, de Gier JW, Daniels R. N-Linked Glycan Sites on the Influenza A Virus Neuraminidase Head Domain Are Required for Efficient Viral Incorporation and Replication. J Virol. 2020:94(19). Epub 2020/07/24. doi: 10.1128/JVI.00874-20. PubMed PMID: 32699088: PubMed Central PMCID: PMCPMC7495393.

20. Yang C, Skiena S, Futcher B, Mueller S, Wimmer E. Deliberate reduction of hemagglutinin and neuraminidase expression of influenza virus leads to an ultraprotective live vaccine in mice. Proc Natl Acad Sci U S A. 2013:110(23):9481–6. Epub 2013/05/22. doi: 10.1073/pnas.1307473110. PubMed PMID: 23690603: PubMed Central PMCID: PMCPMC3677463.

21. Maamary J, Pica N, Belicha-Villanueva A, Chou YY, Krammer F, Gao Q, et al. Attenuated influenza virus construct with enhanced hemagglutinin protein expression. J Virol. 2012:86(10):5782–90. Epub 2012/03/09. doi: 10.1128/JVI.00190-12. PubMed PMID: 22398291: PubMed Central PMCID: PMCPMC3347287.

22. Ramanunninair M, Le J, Onodera S, Fulvini AA, Pokorny BA, Silverman J, et al. Molecular signature of high yield (growth) influenza a virus reassortants prepared as candidate vaccine seeds. PLoS One. 2013:8(6):e65955. Epub 2013/06/19. doi: 10.1371/journal.pone.0065955. PubMed PMID: 23776579: PubMed Central PMCID: PMCPMC3679156.

23. Brooke CB, Ince WL, Wei J, Bennink JR, Yewdell JW. Influenza A virus nucleoprotein selectively decreases neuraminidase gene-segment packaging while enhancing viral fitness and transmissibility. Proc Natl Acad Sci U S A. 2014:111(47):16854–9. Epub 2014/11/12. doi: 10.1073/pnas.1415396111. PubMed PMID: 25385602: PubMed Central PMCID: PMCPMC4250133.

24. Campbell PJ, Danzy S, Kyriakis CS, Deymier MJ, Lowen AC, Steel J. The M segment of the 2009 pandemic influenza virus confers increased neuraminidase activity, filamentous morphology, and efficient contact transmissibility to A/Puerto Rico/8/1934-based reassortant viruses. J Virol. 2014:88(7):3802–14. Epub 2014/01/17. doi: 10.1128/JVI.03607-13. PubMed PMID: 24429367: PubMed Central PMCID: PMCPMC3993553.

25. Plant EP, Ye Z. Chimeric neuraminidase and mutant PB1 gene constellation improves growth and yield of H5N1 vaccine candidate virus. J Gen Virol. 2015:96(Pt 4):752–5. Epub 2014/12/17. doi: 10.1099/jgv.0.000025. PubMed PMID: 25502649.

26. Nordholm J, Petitou J, Ostbye H, da Silva DV, Dou D, Wang H, et al. Translational regulation of viral secretory proteins by the 5' coding regions and a viral RNA-binding protein. J Cell Biol. 2017:216(8):2283–93. doi: 10.1083/jcb.201702102. PubMed PMID: 28696227: PubMed Central PMCID: PMCPMC5551715.

27. Nivitchanyong T, Yongkiettrakul S, Kramyu J, Pannengpetch S, Wanasen N. Enhanced expression of secretable influenza virus neuraminidase in suspension mammalian cells by influenza virus nonstructural protein 1. Journal of virological methods. 2011:178(1-2):44–51. doi: 10.1016/j.jviromet.2011.08.010. PubMed PMID: 21893099.

28. da Silva DV, Nordholm J, Dou D, Wang H, Rossman JS, Daniels R. The influenza virus neuraminidase protein transmembrane and head domains have coevolved. J Virol. 2015:89(2):1094–104. doi: 10.1128/JVI.02005-14. PubMed PMID: 25378494: PubMed Central PMCID: PMC4300628.

29. Wang H, Dou D, Östbye H, Revol R, Daniels R. Structural restrictions for influenza neuraminidase activity promote adaptation and diversification. Nature Microbiology. 2019. doi: 10.1038/s41564-019-0537-z.

30. Kilbourne ED, Schulman JL, Schild GC, Schloer G, Swanson J, Bucher D. Related studies of a recombinant influenza-virus vaccine. I. Derivation and characterization of virus and vaccine. J Infect Dis. 1971:124(5):449–62. Epub 1971/11/01. doi: 10.1093/infdis/124.5.449. PubMed PMID: 5115669.

31. Sanders B, Koldijk M, Schuitemaker H. Inactivated Viral Vaccines. Vaccine Analysis: Strategies, Principles, and Control. 2014:45–80. doi: 10.1007/978-3-662-45024-6_2. PubMed PMID: PMC7189890.

32. Uittenbogaard JP, Zomer B, Hoogerhout P, Metz B. Reactions of beta-propiolactone with nucleobase analogues, nucleosides, and peptides: implications for the inactivation of viruses. J Biol Chem. 2011:286(42):36198–214. Epub 2011/08/27. doi: 10.1074/jbc.M111.279232. PubMed PMID: 21868382: PubMed Central PMCID: PMCPMC3196120.

33. She YM, Cheng K, Farnsworth A, Li X, Cyr TD. Surface modifications of influenza proteins upon virus inactivation by beta-propiolactone. Proteomics. 2013:13(23-24):3537–47. Epub 2013/10/15. doi: 10.1002/pmic.201300096. PubMed PMID: 24123778: PubMed Central PMCID: PMCPMC4265195.

34. Bonnafous P, Nicolai MC, Taveau JC, Chevalier M, Barriere F, Medina J, et al. Treatment of influenza virus with beta-propiolactone alters viral membrane fusion. Biochim Biophys Acta. 2014:1838(1 Pt B):355–63. Epub 2013/10/22. doi: 10.1016/j.bbamem.2013.09.021. PubMed PMID: 24140008.

35. Fan C, Ye X, Ku Z, Kong L, Liu Q, Xu C, et al. Beta-Propiolactone Inactivation of Coxsackievirus A16 Induces Structural Alteration and Surface Modification of Viral Capsids. J Virol. 2017:91(8). Epub 2017/02/06. doi: 10.1128/JVI.00038-17. PubMed PMID: 28148783: PubMed Central PMCID: PMCPMC5375664.

36. da Silva DV, Nordholm J, Madjo U, Pfeiffer A, Daniels R. Assembly of subtype 1 influenza neuraminidase is driven by both the transmembrane and head domains. J Biol Chem. 2013:288(1):644–53. Epub 2012/11/15. doi: M112.424150 [pii] 10.1074/jbc.M112.424150. PubMed PMID: 23150659: PubMed Central PMCID: PMC3537063.

37. Ikematsu H, Kawai N, Tani N, Chong Y, Bando T, Iwaki N, et al. In vitro neuraminidase inhibitory concentration (IC_50_) of four neuraminidase inhibitors in the Japanese 2018-19 season: Comparison with the 2010-11 to 2017-18 seasons. J Infect Chemother. 2020:26(8):775–9. Epub 2020/04/07. doi: 10.1016/j.jiac.2020.03.001. PubMed PMID: 32249161.

38. Chlanda P, Schraidt O, Kummer S, Riches J, Oberwinkler H, Prinz S, et al. Structural Analysis of the Roles of Influenza A Virus Membrane-Associated Proteins in Assembly and Morphology. J Virol. 2015:89(17):8957–66. doi: 10.1128/JVI.00592-15. PubMed PMID: 26085153: PubMed Central PMCID: PMCPMC4524094.

39. Moreno-Pescador G, Florentsen CD, Ostbye H, Sonder SL, Boye TL, Veje EL, et al. Curvature-and Phase-Induced Protein Sorting Quantified in Transfected Cell-Derived Giant Vesicles. ACS Nano. 2019:13(6):6689–701. Epub 2019/06/15. doi: 10.1021/acsnano.9b01052. PubMed PMID: 31199124.

40. Dou D, da Silva DV, Nordholm J, Wang H, Daniels R. Type II transmembrane domain hydrophobicity dictates the cotranslational dependence for inversion. Mol Biol Cell. 2014:25(21):3363–74. doi: 10.1091/mbc.E14-04-0874. PubMed PMID: 25165139.

41. Nordholm J, da Silva DV, Damjanovic J, Dou D, Daniels R. Polar residues and their positional context dictate the transmembrane domain interactions of influenza a neuraminidases. J Biol Chem. 2013:288(15):10652–60. Epub 2013/03/01. doi: M112.440230 [pii] 10.1074/jbc.M112.440230. PubMed PMID: 23447533.

42. Wang N, Glidden EJ, Murphy SR, Pearse BR, Hebert DN. The cotranslational maturation program for the type II membrane glycoprotein influenza neuraminidase. J Biol Chem. 2008:283(49):33826–37. PubMed PMID: 18849342.

43. Saito T, Taylor G, Webster RG. Steps in maturation of influenza A virus neuraminidase. J Virol. 1995:69(8):5011–7. Epub 1995/08/01. PubMed PMID: 7541844: PubMed Central PMCID: PMC189317.

44. Krammer F, Fouchier RAM, Eichelberger MC, Webby RJ, Shaw-Saliba K, Wan H, et al. NAction! How Can Neuraminidase-Based Immunity Contribute to Better Influenza Virus Vaccines? mBio. 2018:9(2). Epub 2018/04/05. doi: 10.1128/mBio.02332-17. PubMed PMID: 29615508: PubMed Central PMCID: PMCPMC5885027.

45. Stuart-Harris CH. A NEUROTROPIC STRAIN OF HUMAN INFLUENZA VIRUS. The Lancet. 1939:233(6027):497–9. doi: https://doi.org/10.1016/S0140-6736(00)74067-0.

46. Appleby JC. The isolation and properties of a modified strain of neurotropic influenza A virus. Br J Exp Pathol. 1952:33(3):280–7. Epub 1952/06/01. PubMed PMID: 14935093: PubMed Central PMCID: PMCPMC2073237.

47. Burnet FM, Lind PE. A genetic approach to variation in influenza viruses; recombination of characters in influenza virus strains used in mixed infections. J Gen Microbiol. 1951:5(1):59–66. Epub 1951/02/01. doi: 10.1099/00221287-5-1-59. PubMed PMID: 14824471.

48. Sugiura A, Ueda M. Neurovirulence of influenza virus in mice. I. Neurovirulence of recombinants between virulent and avirulent virus strains. Virology. 1980:101(2):440–9. Epub 1980/03/01. doi: 10.1016/0042-6822(80)90457-2. PubMed PMID: 361453.

49. Nakajima S, Sugiura A. Neurovirulence of influenza virus in mice. II. Mechanism of virulence as studied in a neuroblastoma cell line. Virology. 1980:101(2):450–7. Epub 1980/03/01. doi: 10.1016/0042-6822(80)90458-4. PubMed PMID: 7361454.

50. Goto H, Kawaoka Y. A novel mechanism for the acquisition of virulence by a human influenza A virus. Proc Natl Acad Sci U S A. 1998:95(17):10224–8. PubMed PMID: 9707628: PubMed Central PMCID: PMC21489.

51. Goto H, Wells K, Takada A, Kawaoka Y. Plasminogen-binding activity of neuraminidase determines the pathogenicity of influenza A virus. J Virol. 2001:75(19):9297–301. doi: 10.1128/JVI.75.19.9297-9301.2001. PubMed PMID: 11533192: PubMed Central PMCID: PMC114497.

52. Schnaar RL, Gerardy-Schahn R, Hildebrandt H. Sialic acids in the brain: gangliosides and polysialic acid in nervous system development, stability, disease, and regeneration. Physiol Rev. 2014:94(2):461–518. Epub 2014/04/03. doi: 10.1152/physrev.00033.2013. PubMed PMID: 24692354: PubMed Central PMCID: PMCPMC4044301.

53. Bourmakina SV, Garcia-Sastre A. Reverse genetics studies on the filamentous morphology of influenza A virus. J Gen Virol. 2003:84(Pt 3):517–27. doi: 10.1099/vir.0.18803-0. PubMed PMID: 12604801.

54. Roberts PC, Lamb RA, Compans RW. The M1 and M2 proteins of influenza A virus are important determinants in filamentous particle formation. Virology. 1998:240(1):127–37. doi: 10.1006/viro.1997.8916. PubMed PMID: 9448697.

55. Calder LJ, Wasilewski S, Berriman JA, Rosenthal PB. Structural organization of a filamentous influenza A virus. Proc Natl Acad Sci U S A. 2010:107(23):10685–90. doi: 10.1073/pnas.1002123107. PubMed PMID: 20498070: PubMed Central PMCID: PMCPMC2890793.

56. Sandbulte MR, Gao J, Straight TM, Eichelberger MC. A miniaturized assay for influenza neuraminidase-inhibiting antibodies utilizing reverse genetics-derived antigens. Influenza Other Respir Viruses. 2009:3(5):233–40. Epub 2009/09/01. doi: 10.1111/j.1750-2659.2009.00094.x. PubMed PMID: 21462400: PubMed Central PMCID: PMCPMC4941552.

57. Hoffmann E, Neumann G, Kawaoka Y, Hobom G, Webster RG. A DNA transfection system for generation of influenza A virus from eight plasmids. Proc Natl Acad Sci U S A. 2000:97(11):6108–13. PubMed PMID: 10801978.

58. Reed LJ, Muench H. A simple method of estimating fifty percent end points. Am J Epidemiol. 1938:27(3):493–7. doi: 10.1093/oxfordjournals.aje.a118408.

59. Couzens L, Gao J, Westgeest K, Sandbulte M, Lugovtsev V, Fouchier R, et al. An optimized enzyme-linked lectin assay to measure influenza A virus neuraminidase inhibition antibody titers in human sera. Journal of virological methods. 2014:210:7–14. doi: 10.1016/j.jviromet.2014.09.003. PubMed PMID: 25233882.

60. Gao J, Couzens L, Eichelberger MC. Measuring Influenza Neuraminidase Inhibition Antibody Titers by Enzyme-linked Lectin Assay. J Vis Exp. 2016:(115). Epub 2016/09/30. doi: 10.3791/54573. PubMed PMID: 27684188: PubMed Central PMCID: PMCPMC5091984.

